# Overexpression of the NEK8 kinase inhibits homologous recombination

**DOI:** 10.1101/2025.02.07.637121

**Authors:** Joshua L. Turner, Georgia Moore, Tyler J. McCraw, Jennifer M. Mason

## Abstract

Homologous recombination maintains genome stability by repairing double strand breaks and protecting replication fork stability. Defects in homologous recombination results in cancer predisposition but can be exploited due to increased sensitivity to certain chemotherapeutics such as PARP inhibitors. The NEK8 kinase has roles in the replication response and homologous recombination. NEK8 is overexpressed in breast cancer, but the impact of NEK8 overexpression on homologous recombination has not been determined. Here, we demonstrate NEK8 overexpression inhibits RAD51 focus formation resulting in a defect in homologous recombination and degradation of stalled replication forks. Importantly, NEK8 overexpression sensitizes cells to the PARP inhibitor, Olaparib. Together, our results suggest NEK8 overexpressing tumors may be recombination-deficient and respond to chemotherapeutics that target defects in recombination such as Olaparib.

## 1. INTRODUCTION

A failure to repair double strand breaks (DSBs) can result in genome instability including genome rearrangements and cell death. Cells repair DSBs through two main pathways. Non-homologous end joining ligates the ends of the DSB back together. Homologous recombination (HR) uses a sister chromatid as a template resulting in repair of the DSB without the loss of genetic information. Inherited mutations in HR genes, including BRCA1 and BRCA2, are often linked to increased cancer risk [1,2]. On the other hand, HR-deficiency sensitizes cells to certain chemotherapeutic drugs including cisplatin and PARP inhibitors such as Olaparib [3,4]. Therefore, understanding how different alterations in cancer impact HR status has important implications for cancer therapy.

Proteins involved in HR have break-dependent and break-independent functions to maintain genome stability. At 5’ resected end of DSBs, RAD51 is loaded onto single-stranded DNA (ssDNA) by mediators including BRCA2 and RAD51 paralogs to form a nucleoprotein filament [5,6]. RAD51 filaments search for a homologous sequence to use as a template for repair. Once found, RAD51 catalyzes the strand exchange reaction by initiating pairing between the ssDNA and the complementary DNA sequence forming an intermediate known as a displacement loop, or D-loop [7,8]. RAD51 is removed from the D-loop structure exposing the 3’OH to allow for DNA synthesis across the region that contains the break. The D-loop is dissolved, and DNA synthesis copies any missing information resulting in repair of the DSB without loss of information. At sites of stalled replication forks, HR proteins have several functions that do not require the presence of a DSB. RAD51 and RAD54 promote fork regression at stalled replication forks [9,10]. RAD51 binds to the regressed fork and is stabilized by several proteins including BRCA2, FANCD2, and FANCA to prevent nuclease-mediated degradation of nascent DNA [11]. Fork protection prevents nuclease degradation and cleavage by nucleases including MRE11, EXO1, DNA2, and MUS81 [12–15].

Homologous recombination is tightly controlled in cells to prevent unscheduled or inappropriate HR. Increased recombination is associated with increased chromosomal rearrangements and has been linked to chemotherapeutic resistance [16]. RAD51 is tightly controlled by proteins that promote nucleoprotein filament assembly and stability including BRCA2 and RAD51 paralogs and disrupt RAD51-DNA interactions including RAD54 and FBH1 [17–20]. RAD51 is also controlled by post-translational modifications that alter RAD51-DNA binding and subcellular localization. RAD51 phosphorylation on residues Ser14 and Tyr13 by PLK1, CK2, and CHK1 promote homologous recombination [21,22]. In contrast, the kinase c-Abl phosphorylates RAD51 on Tyr315 and Tyr54 inhibiting RAD51 nucleofilament formation and nuclear localization [23,24]. Outside of phosphorylation, ubiquitination of RAD51 by FBH1 negatively regulates HR by inhibiting rebinding of RAD51 to DNA [17]. Other HR proteins are also regulated by post translational modifications including RAD54 and BRCA2 [25–27].

Never in mitosis gene A (NIMA) related kinase 8 (NEK8) is a member of a serine/threonine kinase family involved in G2/M progression, DNA repair, centromere localization, and cilia development [28–30]. In primary cilia development, NEK8 regulates cilia length as well as centrosome recruitment [30]. Dominant and recessive mutations in NEK8 leads to juvenile cystic kidney disease (JCK) in mice and nephronophthisis in humans [31,32]. NEK8 plays a role in the G2/M checkpoint by regulating the expression of enzymes such as CDK1 and Cyclin B1 [33]. More recent studies have shown that NEK8 has several roles in S-phase, specifically in DNA repair. NEK8 has been shown to directly interact with ATR in the ATR-CHK1 signaling pathway, regulating the double strand break response and new origin firing [33]. NEK8 promotes RAD51 focus formation at sites of DSBs, HR, and replication fork protection [34].

NEK8 overexpression has been found in a high percentage of non-specific invasive breast cancer [35]. Overexpression of NEK8 in breast cancer carcinomas was found to be involved in increased growth rate, colony formation, and proliferation [36]. Over expression of JCK mutated NEK8 reduced actin expression as well as significantly increase G2/M related protein expression [35]. The role of NEK8 overexpression on tumor progression was associated with its interactions with b-catenin in the Wnt/β-catenin pathway leading to increased cancer cell motility [36]. However, the effects of increased NEK8 expression on homologous recombination have not been explored.

Here, we show that NEK8 overexpression inhibits RAD51 foci formation resulting in a defect in homologous recombination and increased nascent strand degradation of stalled replication forks. Finally, NEK8 overexpression sensitizes cells to the PARP inhibitor, Olaparib. Together, our results indicate NEK8 overexpression may result in HR-defective tumors that may respond to treatment with PARP inhibitors.

## 2. METHODS

### 2.1. Cell culture and drug treatment

hTERT immortalized RPE-1 cells were grown in DMEM/F12 50:50 (Corning, 10-092-CVR) with 10% FBS (Atlas Biologicals, S10350H). HEK293 DR-GFP cells were grown in DMEM (Gibco, 11965–092) with 10% FBS. Cells were incubated at 37°C in 5% CO_2._ Hydroxyurea (Fisher Scientific, AAA1083106) was suspended in water. Olaparib (Thermo Scientific, 466292500) was suspended in DMSO. Cells were treated with the indicated concentrations.

### 2.2. Plasmid transfection

Cells were transfected with MYC-tagged NEK8[32]. Cells were seeded at 200,000 cells in 6-well plates and transfected with Lipofectamine 3000 (Fisher Scientific, L3000001) using manufacturer’s instructions.

### 2.3. siRNA transfections

RPE1 cells were seeded at 100,000 cells in 6-well plates and allowed to adhere for 24 hours. They were then transfected with Lipofectamine RNAiMAX (Fisher Scientific, 13778150) using manufacturers’ instructions. siRNAs against FANCA (Dharmacon, J-019283-07-0020, 75 nM) and RAD51 (Dharmacon, 50 nM) were used. The All-stars negative control (Qiagen) was used as a negative control siRNA. The siRNA sequences are as follows:

siFANCA 5’ GGGCCAUGCUUUCUGAUUU

siRAD51 5’ GACUGCCAGGAUAAAGCUU used in study [37]

### 2.4. Immunofluorescence

Cells were seeded at 40,000 cells in a 12-well plate containing 22 mm circular cover slips. Cells were treated with 2 mM HU or 10 μM Olaparib for 24 hours. Cells were permeabilized (20 mM HEPES, ph7.4, 3 mM MgCl_2_, 50 mM NaCl, 0.5% Triton X-100) for 10 minutes prior to fixation with 3% PFA, 3.4% sucrose. For PCNA staining, cells were fixed with ice-cold methanol. Slides were stained with primary antibodies overnight at 4°C. Primary antibodies used in this study were PCNA (ABCAM, ab18197, 1:1000) and rabbit RAD51 (Pacific Immunology; 1:1000) [37]. Cells were washed 3X with 1X PBS followed by incubation with Alexa Fluor-conjugated secondary antibodies (Invitrogen, 1:1000) for 1 hour at room temperature. Slides were washed 3 times for 5 minutes in 1X PBS followed by 2 minutes each in 70%, 95% and 100% ethanol.

Coverslips were allowed to air dry and mounted with vectashield containing DAPI. Images of 100 PCNA positive nuclei were acquired at 60x magnification on a Zeiss Imager. M2 epifluorescence microscope with an Axiocam 503 mono camera. The number of RAD51 foci in PCNA-positive cells was quantified using Image J software.

### 2.5. Replication fiber analysis

Cells were seeded at 40,000 cells in 12-well plates and allowed to adhere for 24 hours. Cells were sequentially pulsed for 20 minutes with 20 μM CldU and 75 μM IdU. Cells were washed 2X with 1X PBS and treated with 2 mM hydroxyurea for 5 hours before collecting by trypsinization. Untreated controls were collected by trypsinization immediately following the IdU pulse. Cells were resuspended in ice cold 1X PBS at a concentration of 40,000 cells/ml. Replication fibers were prepared as previously described [11]. Slides were stained with mouse BrdU (BD, 347580, 1:50) to label IdU and rat BrdU (Abcam, ab6326, 1:200) to label CldU. Slides were washed with 1X PBS followed by incubation with Alexa Fluor-conjugated antibodies (Invitrogen, 1:1000) Images were acquired at 60x magnification using the Zeiss epifluorescence microscope. The length of at least 150 replication tracts was measured using ImageJ.

### 2.6. Survival assay

Cells were plated in triplicate at 2000 cells per well in black walled 96-well plates (Corning, 3606). Cells were mock treated with DMSO or Olaparib for 4 days before staining with propidium iodide and Hoechst 33342. Live cells were counted and analyzed using the Molecular Devices ImageExpress Micro 4 and proprietary MetaXpress software version 6.7.2.290.

### 2.7. DR-GFP assay

Cells were transfected with empty vector (pCAGG) and I-SCEI+ (pBAS) plasmids [38]. Cells were transfected with siRNA for 24 hours followed by a plasmid transfection for 48 hours as described above. Cells were collected using trypsinization and GFP positive cells were identified by flow cytometry using CytoFLEX and analyzed using FCS Express 7 version 7.16.0047.

### 2.8. Nuclear fractionation

Cells were seeded at 600,000 cells in a 10 cm dish and allowed to adhere for 24 hours. Cells were then treated with 2 mM HU or 10 μM Olaparib for 24 hours. Cells were collected by trypsinization. Nuclear and cytoplasmic fractions were isolated using the nuclear and cytoplasmic extraction kit (Thermo Scientific, 78833) following manufacturer conditions. Protein levels were examined by western blotting.

### 2.9. Western Blotting

Whole cells extracts were prepared by lysing cells (10^6^/100 ml) in Laemmli buffer (62.5mM Tris-HCl, ph6.8, 2% SDS, 10% glycerol, 5% 2-mercaptoethanol, and 0.002% bromophenol blue) and boiled for 10 minutes. Extracts were separated on an SDS-Page gel and transferred to PVDF membrane. Membranes were stained with RAD51 (Novus Biologicals, NBP2-75640, 1:1000), and Vinculin (Cell signaling, 13901S, 1:1000). For the cell fractionation western we stained for Histone H3 (Novus Biologicals, NB500-171, 1:1000) and α-Tubulin (Novus Biologicals, NB100-690, 1:1000). We then stained with HRP secondary antibodies (LI-COR, 926-80011/10, 1:2000) and reacted with Chemiluminescence (LI-COR, 926-95000). Images and analysis were conducted using the LI-COR C-DiGit imager and proprietary Image Studio software version 5.2.5

### 2.10. Statistical analysis

All experiments were performed at least three independent times. Statistical analysis was performed using Graphpad Prism (version 10.2.3). Significance was calculated using an ANOVA followed by Tukey HSD for RAD51 foci, fork deprotection, and DR-GFP assays. To calculate the significance in the survival assay we conducted an ANOVA followed by the Holm-Šídák’s multiple comparisons test. Statistical significance was determined by a p-value below 0.05.

## 3. RESULTS

### 3.1. NEK8 overexpression inhibits RAD51 focus formation

Previous studies have indicated that NEK8 is required for proper RAD51 localization to the sites of double strand breaks [34]. At resected DNA ends, RAD51 forms nucleoprotein filaments that can be visualized by immunofluorescence microscopy. We determined if NEK8 overexpression impacted RAD51 localization by measuring RAD51 focus formation after treatment with hydroxyurea (HU) or the PARP inhibitor, Olaparib (hereafter PARPi) in RPE-1 cells transfected with Myc-tagged NEK8. Cells transfected with an empty vector were used as a control (**Figure 1A**). In control cells, treatment with HU or PARPi significantly increased RAD51 focus formation (**Figure 1B**). Compared to the empty vector control, Myc-NEK8 overexpression resulted in a 2.1-fold decrease in RAD51 focus formation after hydroxyurea and a 1.5-fold decrease in RAD51 focus formation after treatment with PARPi (**Figure 1C**). We did not observe a significant difference in the percentage of PCNA positive cells between control and MYC-NEK8 cells indicating this difference observed between control and NEK8 overexpressing cells is not due to changes in the cell cycle (**Figure S1A**). Together, these results indicate that NEK8 overexpression inhibits RAD51 localization to sites of DNA damage.

**Figure 1.**
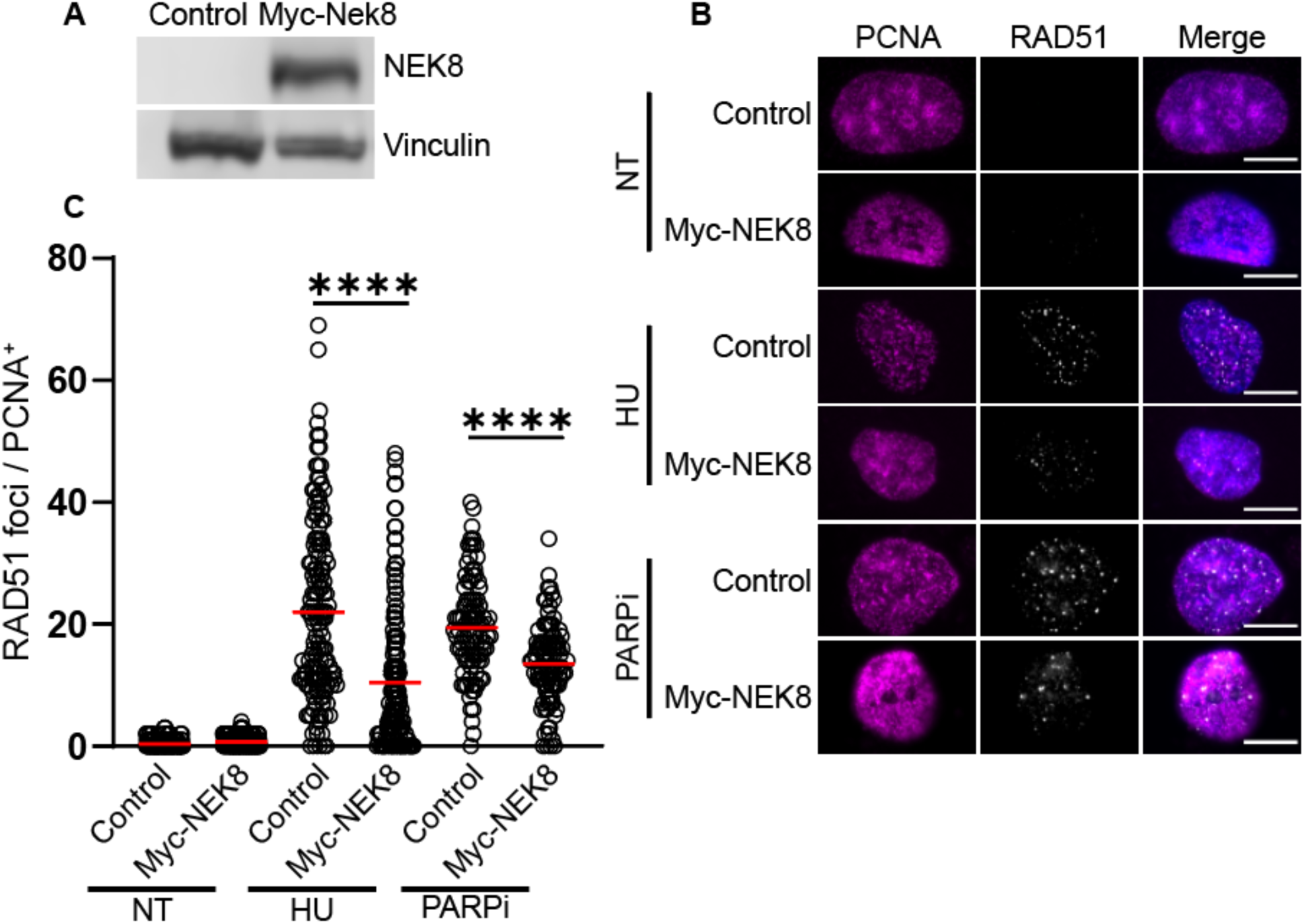
NEK8 overexpression inhibits RAD51 focus formation. **(A)** Western blot showing NEK8 overexpression with vinculin loading control. **(B)** Representative images of PCNA (magenta), RAD51 foci (white), and channels merged with DAPI. Scale bar = 10 μm. **(C)** Dot plot showing RAD51 focus counts for >150 cells per condition, red bar indicates sample mean. N = 3, **** *p*<0.0001 ANOVA; Tukey HSD.

### 3.2. NEK8 overexpression inhibits HR

The reduction of RAD51 focus formation in response to HU and PARPi treatment suggests that NEK8 overexpression may inhibit HR. To test this, we measured HR efficiency using the DR-GFP assay in HEK293 cells [38]. As expected, depletion of RAD51 resulted in a 6.4-fold decrease in HR efficiency (**Figure 2A, B**). Compared to the empty vector control, Myc-NEK8 overexpression resulted in a 3.6-fold decrease in GFP positive cells (**Figure 2B**). These results suggest that the defect in RAD51 localization in NEK8 overexpressing cells is sufficient to inhibit HR.

**Figure 2.**
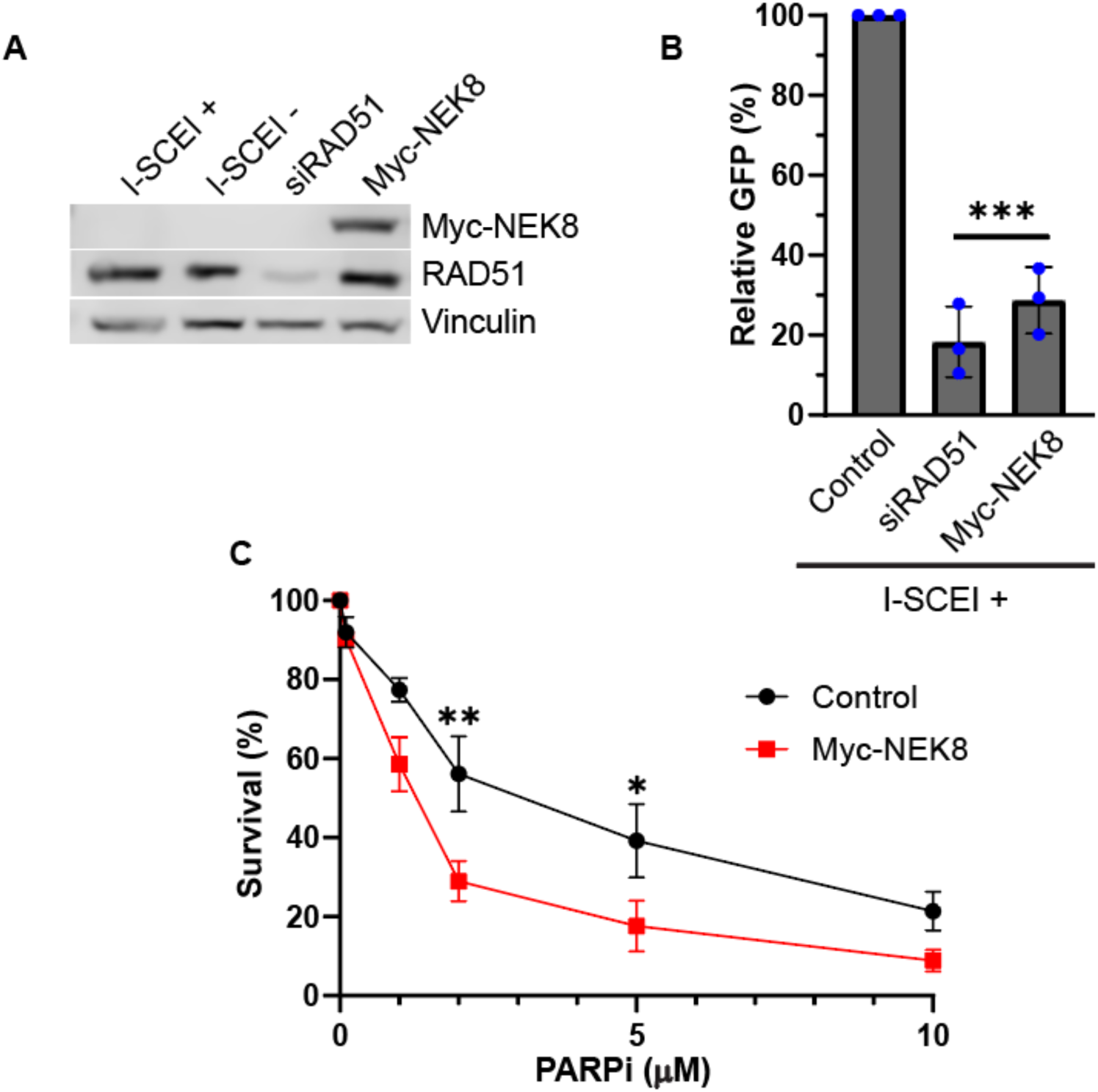
NEK8 overexpression inhibits homologous recombination. **(A)** Western blot showing NEK8 and RAD51 protein levels. Vinculin was used for a loading control. **(B)** Homologous recombination was measured using the DR-GFP assay. Bar graph depicts the percent GFP positive cells relative to the I-SCEI positive control. Individual dots are date points for independent experiments. N = 3, error bars = SD. *p* ***<0.001 ANOVA; Tukey HSD. **(C)** Survival curve of control and Myc-NEK8 overexpressing cells after treatment with indicated PARPi concentrations. N=3, error bars = SEM. \**p*< 0.05, \*\**p*<0.01. ANOVA; Holm-Šídák’s

A defect in HR, such as BRCA2-deficiency, results in increased sensitivity to PARP inhibitors [2]. We tested if MYC-NEK8 over-expression increases sensitivity to PARP inhibitors by treating cells with PARPi. We found that Myc-NEK8 overexpressing cells had decreased survival when treated with PARPi compared to control cells (**Figure 2C**). Together, these results indicate NEK8 overexpression increases sensitivity to PARP inhibition consistent with a defect in HR.

### 3.3. NEK8 overexpression promotes replication fork degradation

At stalled replication forks, RAD51 protects nascent DNA from degradation by nucleases such as MRE11 [39,40]. Therefore, we determined if the decrease in RAD51 localization in NEK8-overexpressing cells results in nascent strand degradation using the replication fiber assay [41]. After sequentially pulsing control and MYC-NEK8 cells with thymidine analogs chlorodeoxyuridine (CldU) and iododeoxyuridine (IdU), cells were treated with hydroxyurea (**Figure 3A**). If replication forks undergo nucleolytic degradation, IdU tracts will be degraded leading to a reduction of the IdU/CldU ratio compared to controls (**Figure 3B,C)**. As a control, we measured replication tract lengths in cells depleted of the fork protection protein, FANCA, and observed a significant decrease in the IdU/CldU ratio (1.04 in controls vs. 0.758 in FANCA-depleted cells) [11]. Myc-NEK8 expressing cells resulted in a significant decrease in IdU/CldU ratio from 1.04 in control cells to 0.747 (**Figure 3C**). These results indicate that overexpressing NEK8 leads to increased nascent strand degradation at stalled replication forks.

**Figure 3.**
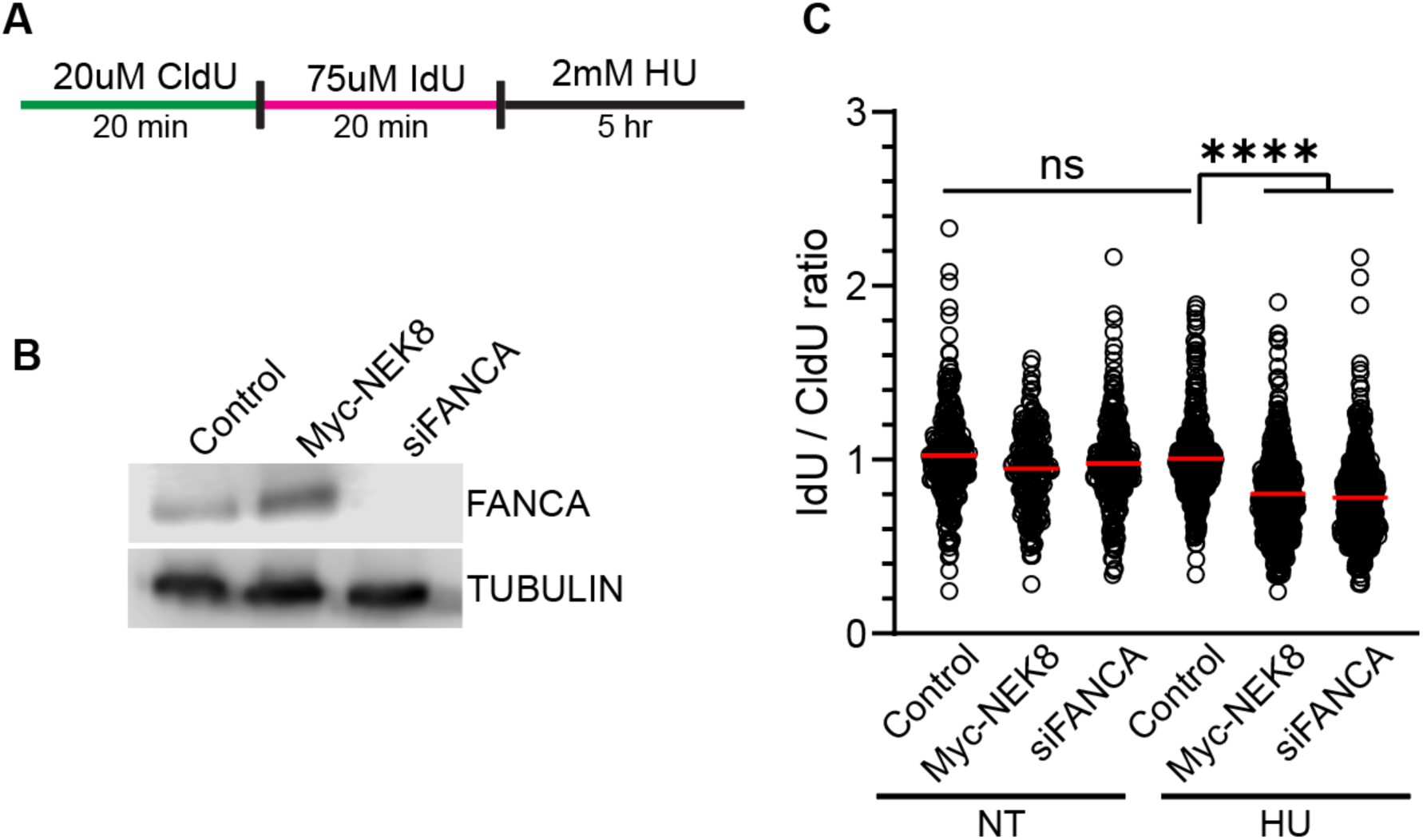
NEK8 overexpression promotes fork deprotection. **(A)** Schematic of experiment used to measure fork deprotection using the replication fiber assay. **(B)** Western blot depicting FANCA protein levels under indicated treatments. Tubulin was used for loading control. **(C)** Dot plot depicting ratio of IdU / CldU tract lengths. Redline represents the mean. >200 fibers per condition, N = 3, ****<0.0001 ANOVA; Tukey HSD.

### 3.4. Changes in RAD51 focus formation is not due to a reduction in nuclear RAD51

It is unclear how NEK8 regulates RAD51 focus formation. It has been shown that post-translational modification of RAD51 can influence its ability to traverse into the nucleus [23]. We determined if loss of RAD51 focus formation is due to a reduction in RAD51 protein levels or decreased RAD51 nuclear localization. To test this, we isolated cytosolic and nuclear protein extracts and analyzed RAD51 expression by western blot. Myc-NEK8 expressing cells showed no significant difference in subcellular localization or steady state levels of RAD51 compared to control cells in untreated cells or after treatment with HU or PARPi **(Figure 4).** This result suggests that impaired RAD51 focus formation in NEK8 overexpressing cells is not due to a reduction in RAD51 localization to the nucleus.

**Figure 4.**
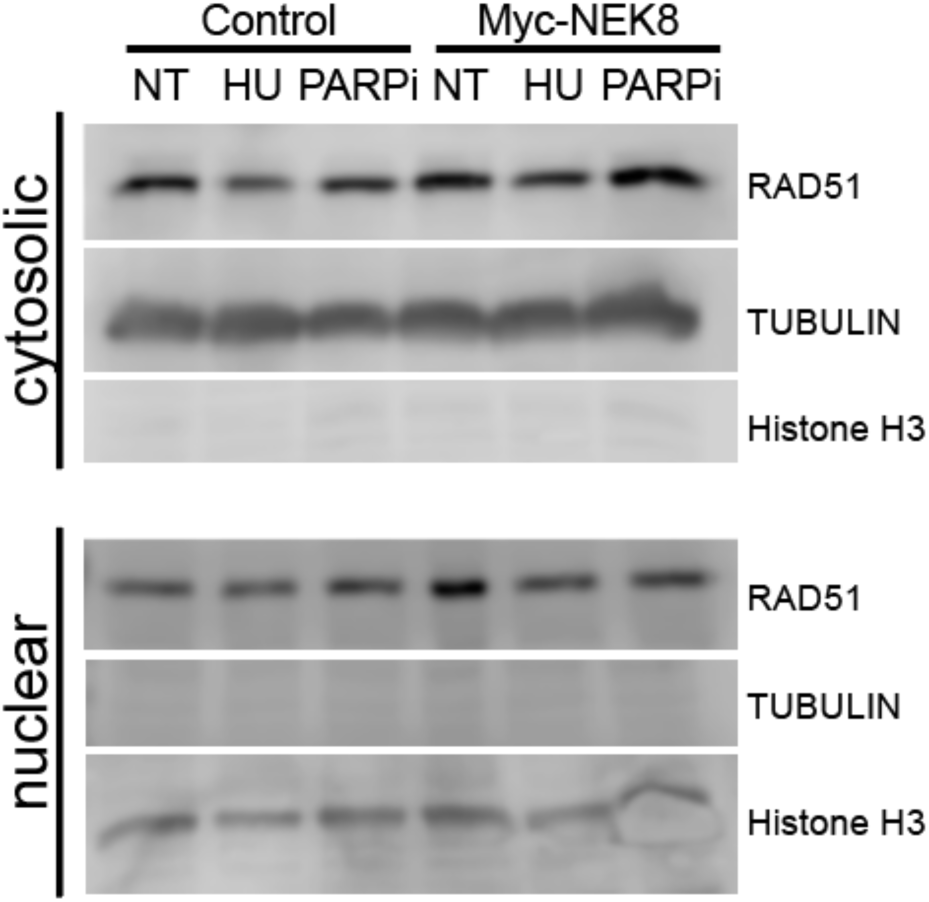
NEK8 overexpression does not impact RAD51 nuclear localization. Western blot depicting RAD51 protein levels in cytosolic (Top) and nuclear fractions (Bottom). Tubulin (cytosolic) + Histone H3 (nuclear) were used as loading controls. N=3

## 4. DISCUSSION

NEK8 expression is essential for RAD51 focus formation and activation of the ATR mediated stress response [34]. However, studying overexpression of NEK8 may provide important clinical relevance due to the prevalence of NEK8 overexpression in invasive breast cancer [36]. In this study, we demonstrated that overexpression of NEK8 caused a significant reduction in RAD51 focus formation after treatment with HU or PARPi. The reduction in RAD51 localization was sufficient to inhibit HR and result in fork deprotection indicating NEK8 overexpression may result in genome instability in tumors.

The mechanism by which NEK8 overexpression inhibits HR was not determined in this study. One possibility is NEK8-mediated phosphorylation of RAD51, or other HR proteins is resulting in reduced RAD51 nucleoprotein filament formation. Phosphorylation of RAD51 at Tyr54 and Tyr315 negatively regulate RAD51 activity [23]. It is unlikely an increase in RAD51 phosphorylation at Tyr54 is leading to a defect in RAD51 focus formation in NEK8 overexpressing cells. Phosphorylation of RAD51 at Try54 inhibits the import of RAD51 to the nucleus and we did not observe a significant difference in RAD51 protein levels in the nuclear fraction in NEK8 overexpressing cells [23]. Another possibility is NEK8 is leading to increased phosphorylation of other HR proteins such as BRCA2. For instance, CDK phosphorylates BRCA2 at Ser3291 to disrupt the interaction between the C-terminal region of BRCA2 and RAD51[25]. NEK8 regulates Cyclin A-associated CDK activity, but it is not clear if NEK8 has roles in other CDK-dependent functions [33]. Future work will need to be conducted to determine how NEK8 regulates HR and if regulation occurs through direct phosphorylation of HR proteins.

NEK8 overexpression plays a direct role in the proliferation and invasiveness of many breast cancers [36]. Knocking down NEK8 was effective in reducing tumor proliferation but loss of NEK8 also reduced RAD51 localization, promoting HR deficiency [34,36]. We demonstrate that NEK8 overexpression also results in a decrease in RAD51 focus formation and HR deficiency. Cancer cells defective in HR (e.g. BRCA2 deficient breast cancer) are hypersensitive to PARP inhibitors such as Olaparib [42]. Consistent with a defect in HR, we found NEK8 overexpression increased sensitivity to Olaparib. Thus, non-specific breast cancers containing increased NEK8 expression may be sensitive to PARP inhibitors. Future work will need to be conducted to determine how breast cancer cells with elevated NEK8 respond to PARP inhibition.

---

**Figure S1.**
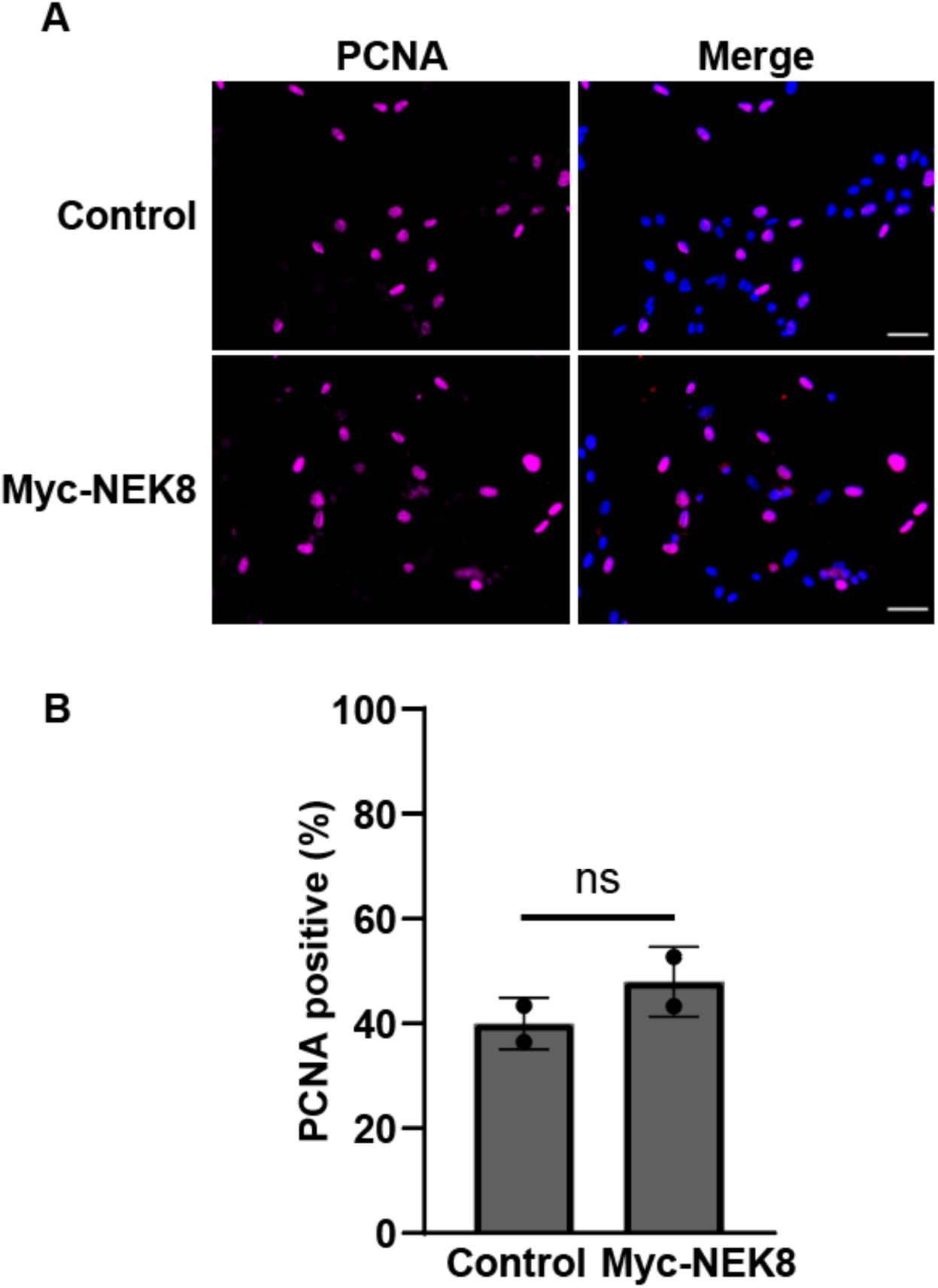
NEK8 overexpression does not alter the percentage cells in S-phase. **(A)** Representative images of PCNA (magenta) and DAPI (Blue), scale bar = 50μm. **(B)** Bar graph of PCNA positive cells. Dots represent individual data points from independent experiments. N=2. ANOVA; Tukey HSD.

**Figure S2.**
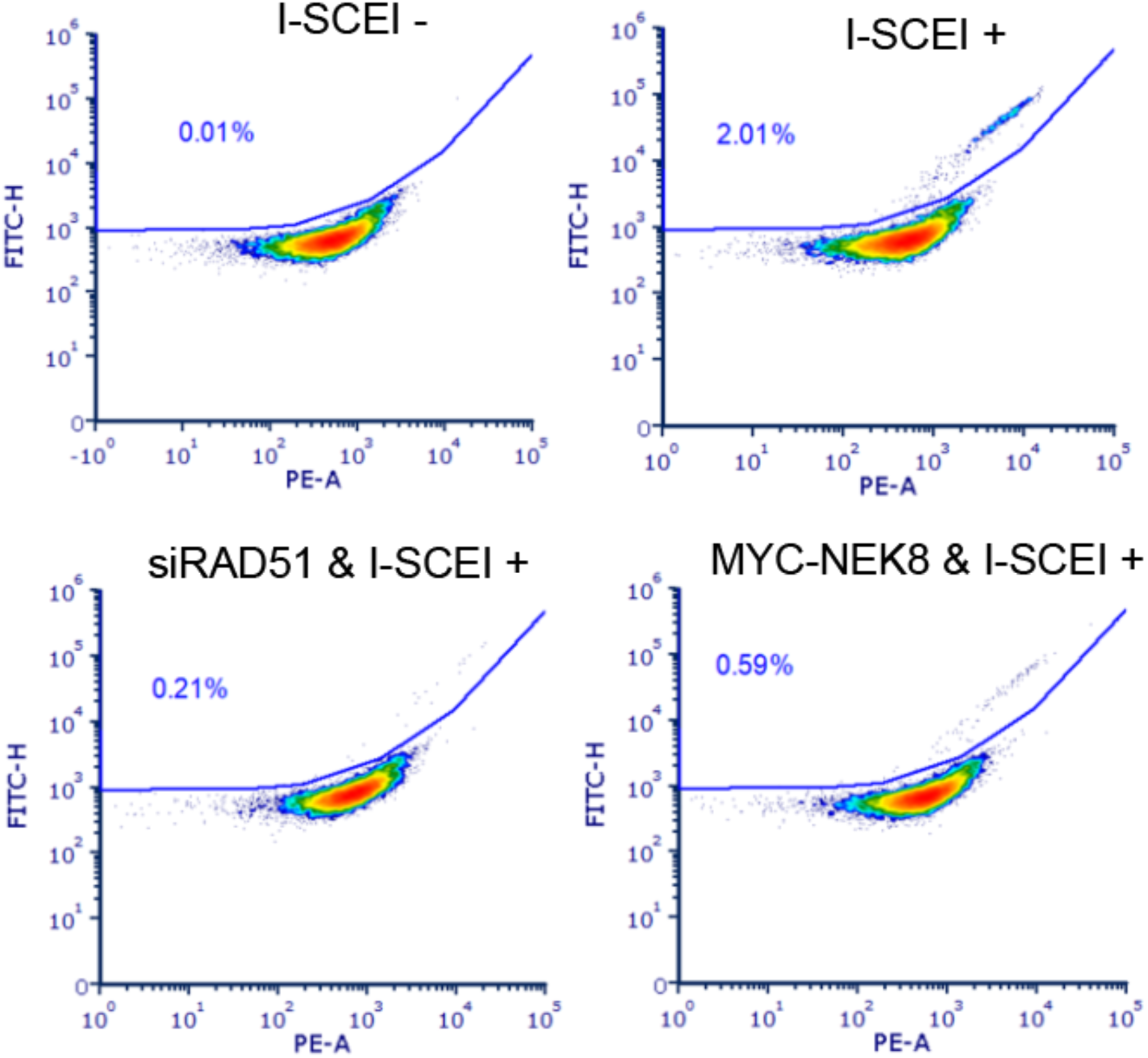
DR-GFP assay to measure HR (HR+). Representative intensity plots depicting HR positive (GFP+) cells in the indicated samples. X-axis is PE, y-axis is GFP. Gates defining GFP+ population is shown in blue. Quantification of assay is in Figure 2B.

